# Aberrant neuronal hyperactivation causes an age- and diet-dependent decline in associative learning behavior

**DOI:** 10.1101/2024.03.21.586045

**Authors:** Binta Maria Aleogho, Mizuho Mohri, Moon Sun Jang, Sachio Tsukada, Yana Al-Hebri, Yuki Tsukada, Ikue Mori, Kentaro Noma

**Affiliations:** Group of Microbial Motility, Department of Biological Science, Division of Natural Science, Graduate School of Science, Nagoya University, Nagoya, Japan; Group of Nutritional Neuroscience, Neuroscience Institute, Graduate School of Science, Nagoya University, Nagoya, Japan; Group of Molecular Neurobiology, Neuroscience Institute, Graduate School of Science, Nagoya University, Nagoya, Japan; Milk Science Research Institute, MEGMILK SNOW BRAND Co. Ltd., Saitama, Japan

## Abstract

The impairment of memory, cognition, and behavior during aging is generally thought to arise from diminished neuronal activities. The nematode *Caenorhabditis elegans* exhibits age-dependent declines in an associative learning behavior called thermotaxis. Genetic ablation of individual neurons revealed that an absence of either AWC sensory or AIA inter-neurons preserved the thermotaxis ability of aged animals. Calcium imaging showed age-dependent spontaneous hyperactivities in both neurons. The age-dependent neuronal hyperactivity and behavioral decline were ameliorated by changing diets. We further demonstrate that the enhanced activities of AWC and AIA were differentially dependent on the forms of neurotransmission mediated by neurotransmitters and neuropeptides. Together, our data provides evidence that aberrantly enhanced, not diminished, neuronal responses can impair behavior during aging.

**One-Sentence Summary:** Enhanced neuronal activity during aging impairs *C. elegans* learning behavior.

## Main Text

Aging presents a broad spectrum of systemic alterations in animals that can impair an animal’s sensory perception and behavioral functions. Age-dependent sensory impairment, memory loss, and cognitive and behavioral decline are generally attributed to neuron loss, synaptic dysfunction, and decreased neuronal activities over time (*1, 2*). At the same time, neuronal hyperactivity is reported in humans and other organisms during aging (*3, 4*). However, it is unclear whether neuronal hyperactivity is the cause of cognitive impairment or a compensatory mechanism of circuit dysfunction. It is crucial to examine age-dependent changes in neuronal activities and clarify their consequences on cognitive function.

The roundworm *Caenorhabditis elegans* (*C. elegans*) with a simple nervous system of only 302 neurons, which are mostly in a predetermined configuration, is ideal for studying the causal relationship between neuronal activities and behavioral consequences. *C. elegans* exhibits thermotaxis, in which animals associate food availability with their cultivation temperature (T_cult_) and migrate toward T_cult_ on a thermal gradient (*5-7*). Young *C. elegans* fed *Escherichia coli (E. coli*) as a standard laboratory diet can perform thermotaxis, while aged animals show a decline in this behavior (*8, 9*). We reported that animals fed another diet, *Lactobacillus reuteri* (*L. reuteri*), can preserve thermotaxis ability even when aged (*9*). The *C. elegans* thermotaxis circuit and its molecular and cellular mechanisms have been well characterized in young animals (*6, 10-16*). Much less, however, is known about the neuronal basis for the age- and diet-dependent thermotaxis decline.

### Aged animals require AFD sensory and AIY inter-neurons for thermotaxis

AFD and AIY are the major sensory and inter-neurons, respectively, required for thermotaxis in young (Day 1 of adulthood) animals (*6, 11, 17, 18*) (Fig. 1A). We asked if these neurons were also required in aged (Day 5 of adulthood) animals fed different diets (Fig. 1B). Genetic ablation of AFD or AIY neurons dramatically reduced the thermotaxis performance index (Fig. 1, C, D, and F), rendering animals athermotactic (Fig. 1E) and cryophilic (Fig. 1G), respectively, regardless of age or diet. We reasoned that different activities of those neurons might underlie the age- and diet-dependent changes in thermotaxis and examined their temperature-evoked calcium responses. To this end, we simultaneously recorded RCaMP2 signals in AFD and AIY (Fig. 1H) upon a shallow temperature rise from 21.5 to 22.5°C (0.05°C/s). However, no drastic changes were observed across all ages and diets (Fig. 1, I to L). Imaging only AFD using a steeper temperature rise from 19 to 24°C (fig. S1A) also revealed no apparent difference (fig. S1, B to E). The combined results demonstrate that animals require both AFD and AIY neurons for thermotaxis behavior, but that the age- and diet-dependent thermotaxis decline cannot be explained by changes in the activities of these neurons.

**Fig. 1.**
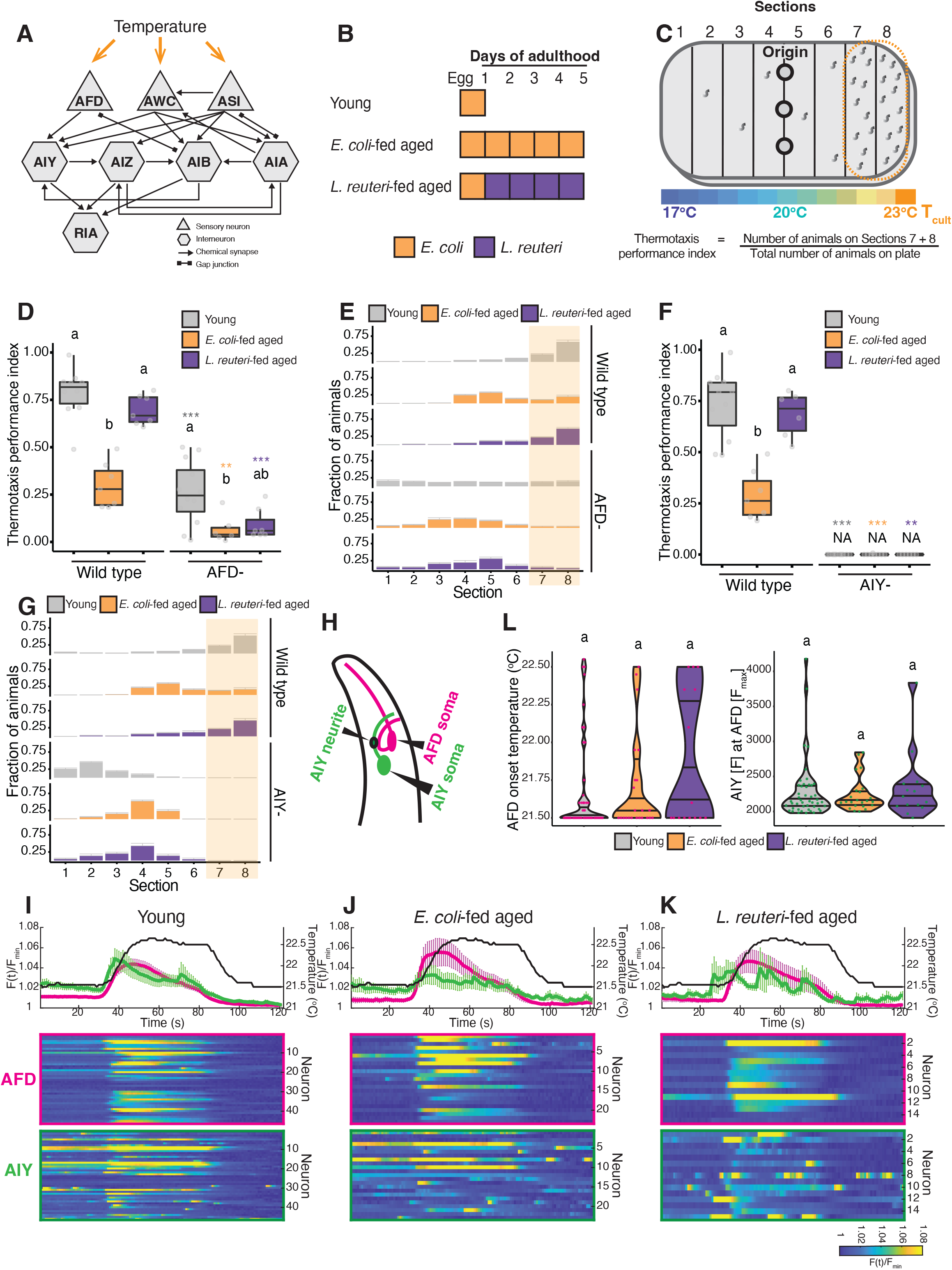
Aged animals require AFD sensory and AIY inter-neurons for thermotaxis. **(A)** Schematic of the neural circuit diagram of thermosensory and inter-neurons (22, 32-34). **(B)** Schematic of animal feeding design. “Young” refers to animals at the first day of adulthood (D1), while “aged” refers to animals at the fifth day of adulthood (D5). **(C)** Schematic of a thermotaxis assay plate. Eight sections were drawn on the plate, and animals were spotted at the origin (20°C center, except otherwise stated) and allowed to move freely. Thermotaxis performance indices reflecting the animals’ ability to learn and migrate to T_cult_ were calculated using the indicated formula. **(D**,**F)** Box and whisker plots of thermotaxis performance indices of wild type and indicated neuron-ablated animals in the different age and diet conditions. ‘Not applicable’ (NA) in (F) depicts that the most values were zero and therefore could not be analyzed statistically. **(E**,**G)** Distribution of animals with indicated genotypes on thermotaxis plates from data in (D,F) respectively. Light brown rectangles indicate the two sections near T_cult_. Error bars denote SEM. **(H)** Schematic of AFD thermosensory neuron and AIY interneuron. **(I**,**J**,**K)** Calcium signals of AFD soma and AIY neurite in indicated conditions in the wild type under a 1°C temperature increase at 0.05°C/s. RCaMP2 fluorescence, F(t), in AFD and AIY was simultaneously measured and standardized by the minimum fluorescence value, F_min_, in each neuron recording. Magenta and green lines depict the average calcium signals in AFD and AIY, respectively. Error bars are SEM. Black lines indicate average temperature stimuli. Heat maps of calcium signals in AFD and AIY are shown at the bottom. **(L)** Violin plots showing temperature at the onset of AFD activation (left) and the AIY fluorescence value, AIY [F], corresponding to AFD maximum fluorescence, AFD [F_max_], (right) from data in (I,J,K). Middle lines within violin plots show medians, and top and bottom lines show interquartile ranges. Each data point is data obtained from a single neuron. Statistical analysis was by Kruskal Wallis test followed by post-hoc Steel-Dwass test for comparisons within a genotype. Different alphabets depict significant difference. Post-hoc Steel test was used for corresponding comparisons with control within a condition, as represented by the distinct asterisk colors. **p ≤0.01 and ***p ≤ 0.001.

### Loss of AWC sensory and AIA inter-neurons preserved thermotaxis ability in aged animals

We then asked whether other neurons playing a role in thermotaxis behavior in young *C. elegans* (*6, 11, 17, 19-22*) (Fig. 1A) contribute to thermotaxis in aged animals. We individually ablated AWC and ASI sensory neurons and AIZ, AIB, AIA, and RIA interneurons by cell-specifically expressing the reconstituted caspase during development (*23*). Ablation of ASI, AIZ, AIB, or RIA did not change the thermotaxis performance of aged animals irrespective of diets. Surprisingly, individually ablating AWC or AIA maintained, instead of impaired, thermotaxis in *E. coli-*fed aged animals (Fig. 2A). Since ablating either neuron did not drastically alter thermotaxis in young and *L. reuteri*-fed animals (Fig. 2A), it is plausible that AWC and AIA are dispensable for thermotaxis. We eliminated the possibility that ablating AWC and AIA neurons caused *E. coli*-fed aged animals to be thermophilic by performing thermotaxis by spotting animals on a 23°C, instead of the usual 20°C, center (fig. S2, A and B). These observations imply that AWC and AIA impair thermotaxis in *E. coli*-fed aged animals.

**Fig. 2.**
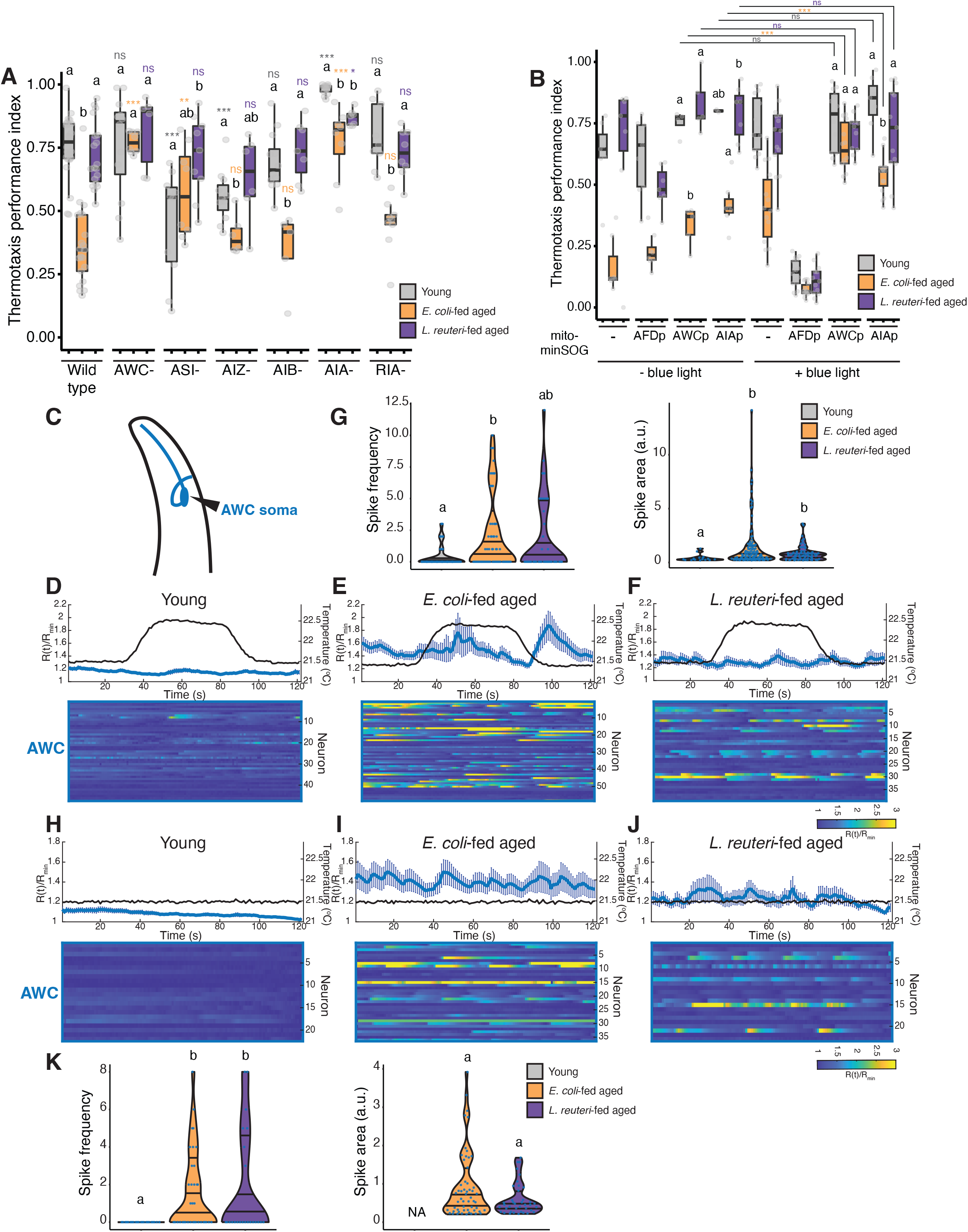
Hyperactivity of AWC sensory neuron interferes with thermotaxis in aged animals. **(A**,**B)** Box and whisker plots of thermotaxis performance indices of wild type and animals with indicated neuron ablation in the different diet and age conditions. **(A)** Cell ablation by caspase during development. **(B)** Cell ablation by mito-miniSOG one day prior to the thermotaxis assay. **(C)** Schematic of AWC sensory neuron. **(D**,**E**,**F)** Calcium signals of AWC soma in indicated conditions in the wild type in response to a 1°C temperature increase at 0.05°C/s. GCaMP6f in AWC and TagRFP in AWC^ON^ (used as a reference) were measured. The GCaMP6f/TagRFP ratio, R(t), was calculated and standardized by the minimum fluorescence ratio, R_min_, in each neuron recording. Blue lines depict the average calcium signals in AWC. Error bars are SEM. Black lines indicate average temperature stimuli. Heat maps of calcium signals are shown at the bottom. **(G)** Violin plots showing quantification of calcium spike frequency per soma (left) and calcium spike area per soma (right) from data in (D,E,F). Each data point represents one soma. **(H**,**I**,**J)** Calcium signals of AWC soma in indicated conditions in wild type under a constant temperature stimulus. Genotype, measurement standardization, and plots and heat map descriptions are same as in (D,E,F). **(K)** Violin plots showing quantification of calcium spike frequency per soma (left) and calcium spike area per soma (right) from data in (H,I,J). ‘Not applicable’ (NA) in the right plot depicts that no areas could be measured since no spikes were detected in all neurons recorded (left). Each data point represents one soma. Statistical analysis was by Kruskal Wallis test followed by post-hoc Steel-Dwass test for comparisons within a genotype. Different alphabets depict significant difference. Post-hoc Steel test was used for corresponding comparisons with control within a condition, as represented by the distinct asterisk colors. *p ≤ 0.05, **p ≤ 0.01, ***p ≤ 0.001, and ‘ns’ indicates not significant (p>0.05).

Our cell ablation experiments by the reconstituted caspase indicate that the absence of AWC or AIA during development, aging, and/or behavioral assays is required to maintain thermotaxis in aged animals. To address the timing of the AWC and AIA ablations, we ablated specific neurons one day before the assay by illuminating blue light on strains expressing mitochondria-targeted mini Singlet Oxygen Generator (mito-miniSOG) (*15, 24*). As a control, animals with AFD thermosensory neurons ablated by mito-miniSOG exhibited low thermotaxis performance irrespective of age and diet (Fig. 2B). Ablation of AWC neurons one day prior to the assay preserved thermotaxis of *E. coli*-fed aged animals (Fig. 2B) similarly to the cell ablation by caspases (Fig. 2A), suggesting that AWC probably impaired thermotaxis during the assay and not during development or aging. Ablation of AIA neurons significantly increased the thermotaxis performance of *E. coli*-fed aged animals but did not restore it to the level of no blue light condition (Fig. 2B).

### AWC sensory neurons are hyperactive in *E. coli*-fed aged animals

Given that AWC-ablated animals showed improved thermotaxis performance in the *E. coli*-fed aged condition, we reasoned that AWC sensory neurons likely disturb the thermosensory circuit in these animals. AWC can respond to strong temperature stimuli as thermosensory neurons (*11, 21*) and show feeding state-dependent modulation in young animals (*25*). Therefore, we performed calcium imaging of AWC (Fig. 2C) to investigate responses to a shallow temperature rise from 21.5 to 22.5°C (0.05°C/s). Young animals showed minimal to no temperature-evoked responses (Fig. 2D), whereas aged animals displayed striking stochastic hyperactive responses (Fig. 2, E and F), with the *E. coli*-fed aged group showing significantly more calcium spikes than the *L. reuteri*-fed aged group (Fig. 2G). The calcium spike areas of both aged groups, while higher than those in young animals, were comparable (Fig. 2G). To check the temperature dependency of these stochastic responses, we imaged animals at a constant temperature of 21.5°C. Similar to the temperature-evoked context, young animals did not show spikes (Fig. 2H), whereas the stochastic hyperactivities remained in aged animals irrespective of diets (Fig. 2, I and J). Both aged groups showed comparable spike frequencies and spike areas (Fig. 2K), although *L. reuteri*-fed animals in the temperature-evoked and temperature-independent contexts appeared to have lower activities than their *E. coli*-fed counterparts (Fig. 2, F and J). These results imply that aging causes an increased frequency of spontaneous AWC activities, and in combination with the behavioral results of AWC-ablated animals (Fig. 2, A and B), we speculate that AWC hyperactivity in aged animals disrupts thermotaxis behavior.

### Hyperactivity of AWC causes thermotaxis defects irrespective of age or diet

To address the hypothesis that hyperactive AWC sensory neurons disrupt thermotaxis behavior in aged animals, we tested a mutant with AWC hyperactivity caused by a loss of function of *srtx-1* gene encoding a G-protein-coupled receptor (*21, 26*). Young *srtx-1* mutants displayed thermotaxis defects (fig. S3), consistent with previously reported defects in isothermal tracking, which is the animals’ ability to continue along the same temperature on a thermal gradient (*21*). Moreover, both *E. coli*-fed and *L. reuteri*-fed aged *srtx-1* mutants also exhibited thermotaxis defects (fig. S3). These defects were at least partially dependent on the AWC because ablation of AWC significantly suppressed the defects (fig. S3). These results suggest that AWC hyperactivity disrupts the thermotaxis circuit to impair the thermotaxis behavior.

### AWC hyperactivity in aged animals involves neuronal communication

To assess whether the hyperactivities of AWC in *E. coli*-fed aged animals occurred cell-autonomously or not, we performed AWC calcium imaging in the mutants of two genes required for neuronal communication: *unc-13* encoding a presynaptic protein required for synaptic vesicle docking, priming, and fusion for neurotransmitter release (*27-29*); *unc-31* encoding a protein required for dense-core vesicle docking, priming, and exocytosis in neuropeptide secretion (*28, 30, 31*). In the *unc-13* mutants, AWC hyperactivities in *E. coli*-fed aged animals were abolished, as reflected by the significantly lower spike frequency than the wild type (Fig. 3, A and B). Only one outlier among 20 individuals in the *E. coli*-fed aged *unc-13* mutants accounted for the higher spike area observed. There was no significant difference in either spike frequency or area between the *L. reuteri*-fed aged *unc-13* mutants and wild-type animals (Fig. 3B). These results suggest that AWC hyperactivity requires synaptic transmission from other neurons. By contrast, in *unc-31* mutants, *E. coli*-fed aged animals continued to show stochastic hyperactivity, whereas *L. reuteri*-fed aged animals showed little to no responses (Fig. 3C). The spike frequencies of both aged groups were comparable to the wild type (Fig. 3D). These results suggest that AWC hyperactivity in aged animals is independent of *unc-31*. We note that the spike area of *E. coli*-fed aged *unc-31* animals is significantly less than the wild-type counterpart, suggesting that neuropeptides might play a role in amplifying signals. These findings suggest that upstream neuron(s) regulate AWC hyperactivation mainly through *unc-13*-mediated neurotransmitter release.

**Fig. 3.**
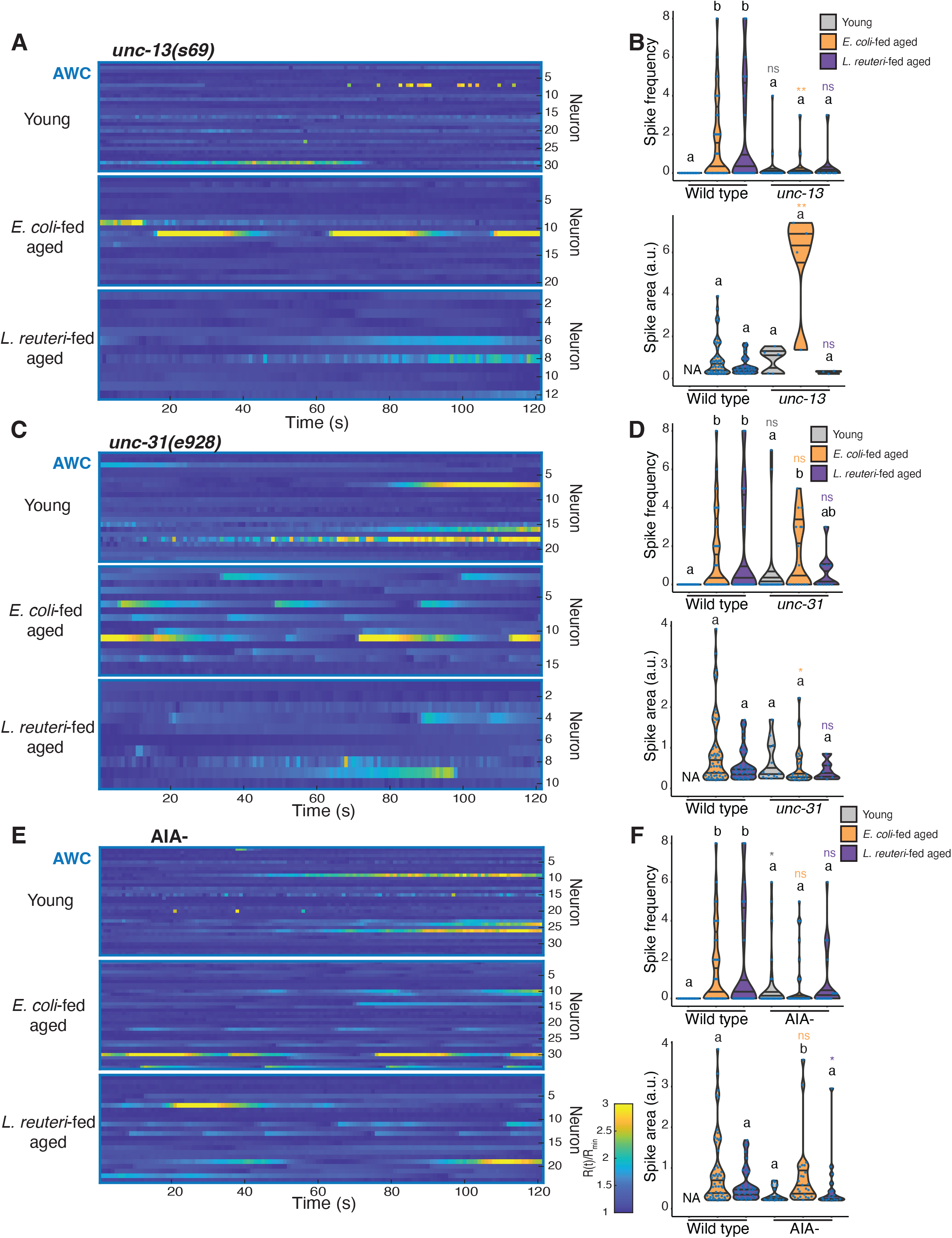
AWC hyperactivity in aged animals is dependent on *unc-13* and partially on AIA interneuron. **(A**,**C**,**E)** Heat maps of calcium signals of AWC soma in indicated conditions in *unc-13(s69*) mutants, which disrupt synaptic vesicle transmission, *unc-31(e928*) mutants, which disrupt neuropeptide transmission, and AIA-ablated animals, respectively under a constant temperature stimulus of 21.5°C. **(B**,**D**,**F)** Quantification of calcium spike frequency per soma (left) and calcium spike area per soma (right) from data in (A,C,E) respectively. Wild type data are the same as in **Fig. 2K**. Statistical analysis was by Kruskal Wallis test followed by post-hoc Steel-Dwass test for comparisons within a genotype. Different alphabets depict significant difference. Post-hoc Steel test was used for corresponding comparisons with control within a condition, as represented by the distinct asterisk colors. *p ≤ 0.05, **p ≤ 0.01, and ‘ns’ indicates not significant (p>0.05).test was used for corresponding comparisons with control within a condition, as represented by the distinct asterisk colors. *p ≤ 0.05, **p ≤ 0.01, ***p ≤ 0.001, and ‘ns’ indicates not significant (p>0.05).

Information from *C. elegans* neural networks and connectome analysis (*32-34*) (http://wormweb.org/) show that AWC is post-synaptic to several sensory and inter-neurons, including AIA interneurons. Given that the genetic ablation of AIA phenocopied that of AWC (Fig. 2A), we asked if AIA is the source of AWC hyperactivity. To test this, we performed AWC calcium imaging in an AIA-ablated strain. Aged animals, irrespective of diets, showed diminished AWC hyperactivities compared to the wild type, although not statistically significant (Fig. 3, E and F). This observation suggests a contribution of AIA to AWC hyperactivity.

### AIA interneurons are hyperactive in aged animals

If AIA contributes to AWC hyperactivity, the AIA interneurons are also likely to show hyperactivities during aging. Therefore, we imaged the neurite of AIA interneurons (Fig. 4A) at a constant temperature of 21.5°C and revealed that *E. coli-*fed aged animals showed a significantly higher spike frequency and larger spike area than young and *L. reuteri*-fed aged animals (Fig. 4, B to E). These results suggest that, like AWC sensory neurons, AIA interneurons are spontaneously hyperactive in *E. coli-*fed aged animals. In combination with behavioral results of AIA ablation (Fig. 2A,B), hyperactivity of AIA neurons seem to impair thermotaxis behavior in aged animals.

**Fig. 4.**
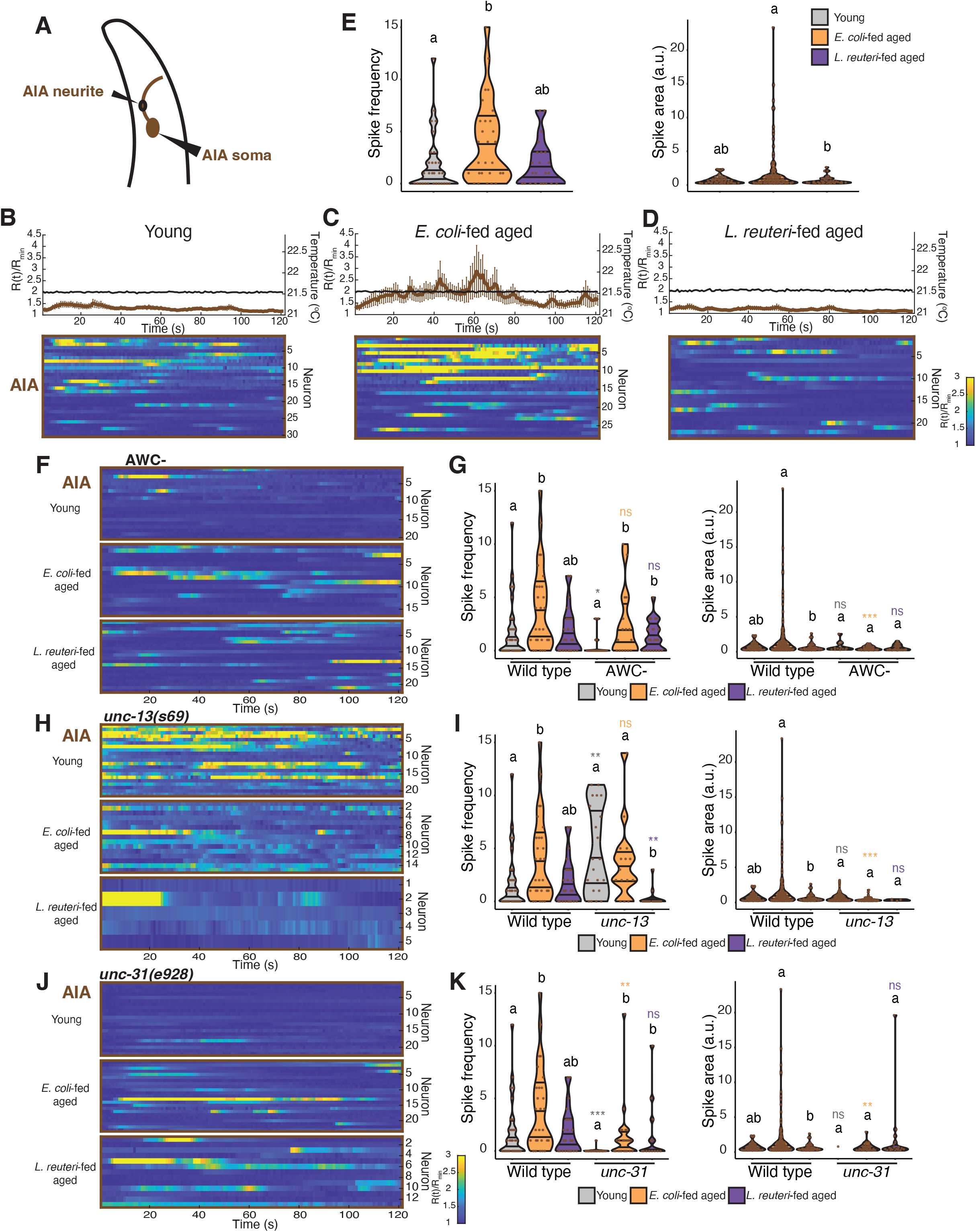
AIA interneuron exhibits age-dependent hyperactivity that depends on *unc-31* and partially on AWC. **(A)** Schematic of AIA interneuron. **(B**,**C**,**D)** Calcium signals of AIA neurite in indicated conditions in the wild type under a constant temperature stimulus. YFP/CFP fluorescence ratio, R(t), of AIA was measured and standardized by the minimum fluorescence ratio, R_min_, in each neuron recording. Brown lines depict the average calcium signals in AIA. Error bars are SEM. Black lines indicate average temperature stimuli. Heat maps of calcium signals are shown at the bottom. **(E)** Violin plots showing quantification of calcium spike frequency per soma (left) and calcium spike area per soma (right) from data in (B,C,D). Each data point represents one soma. **(F**,**H**,**J)** Heat maps of calcium signals of AIA neurite in indicated conditions in AWC-ablated animals, *unc-13* mutants, and *unc-31* mutants, respectively under a constant temperature stimulus. Measurements standardization is same as in (B,C,D). **(G**,**I**,**K)** Quantification of calcium spike frequency per soma (left) and calcium spike area per soma (right) in (F,H,J). Wild-type data are the same as in (E). Statistical analysis was by Kruskal Wallis test followed by post-hoc Steel-Dwass test for comparisons within a genotype. Different alphabets depict significant difference. Post-hoc Steel test was used for corresponding comparisons with control within a condition, as represented by the distinct asterisk colors. *p ≤ 0.05, **p ≤ 0.01, ***p ≤ 0.001, and ‘ns’ indicates not significant (p>0.05).

AWC and AIA neurons share bidirectional synapses (*34*). To evaluate a possible AWC contribution to AIA hyperactivity, we imaged AIA interneurons in an AWC-ablated background. AWC ablation diminished the hyperactive response of AIA in the *E. coli-*fed aged group (Fig. 4F) with less spike area, although the frequency was not significantly reduced (Fig. 4G). This observation implies that AWC neurons partially contribute to AIA hyperactivity. These results also do not undermine the possible contribution from a shared upstream neuron of both AWC and AIA neurons.

Asking which group of neuromodulators might regulate AIA interneuronal communication, we imaged AIA neurons in *unc-13* and *unc-31* mutant backgrounds. In all three groups of *unc-13* mutants, calcium spikes were fragmented and transient and occurred frequently within individuals (Fig. 4H). Spikes in young animals occurred more frequently than in the wild type, whereas those in *E. coli*-fed aged animals remained largely unchanged (Fig. 4, H and I). These results suggest that AIA neurons receive an inhibitory input in young animals, as in the context of olfaction (*35*), but do not in aged animals. Calcium dynamics in *unc-31* mutants showed a drastic reduction in AIA activity in young and *E. coli*-fed aged animals but not in *L. reuteri*-fed aged animals (Fig. 4, J and K). The spike area of *E. coli*-fed aged animals was also significantly lower than in the wild type (Fig. 4K), suggesting that *unc-31*-mediated neuropeptide signaling plays a role in regulating AIA hyperactivity in young and aged animals.

### *daf-16* functions in AWC neurons to maintain thermotaxis in *L. reuteri*-fed aged animals

We reported that the *C. elegans* ortholog of the mammalian FOXO transcription factor (*36, 37*), DAF-16, involved in various biological processes (*38*), plays a role in the amelioration of age-dependent thermotaxis decline by feeding *L. reuteri* (*9*). This *L. reuteri*-dependent ameliorative effect was dependent on *daf-16* b isoform functioning in neurons (*9*). However, the responsible neurons were left unidentified. Since we now revealed that AWC and AIA hyperactivation impaired the thermotaxis behavior, we asked if *daf-16b* functions in these neurons. We found that *L. reuteri*-fed aged animals expressing *daf-16b* in AWC neurons, but not in AIA neurons, rescued the *daf-16* phenotype (fig. S4). Imaging AWC neurons of *daf-16(mu86*) mutants revealed that the *L. reuteri*-fed aged group showed hyperactivity with significantly more spikes than the other two groups (fig. S5). These results suggest that *daf-16* functioning in AWC is required for *L. reuteri* to prevent age-dependent thermotaxis decline.

### AWC and AIA neuronal hyperactivity is independent of olfaction

AWC and AIA neurons are required for chemotaxis to volatile odorants such as benzaldehyde, isoamyl alcohol, and 2-butanone (*39-41*). To ask whether the age-dependent hyperactivity acts through olfaction, we performed thermotaxis on three mutants selected for their defective chemotaxis to AWC-sensed odorants: *odr-1(n1936), odr-2(n2145*), and *odr-5(ky9*) (*42*). In the *E. coli*-fed condition, *odr-1* and *odr-5* mutants exhibited thermotaxis decline as in the wild type (fig. S6), suggesting that AWC hyperactivity is independent of its olfactory function. On the other hand, the *odr-2* mutants showed better thermotaxis ability than the wild type across ages and diets (fig. S6). This result suggests that *odr-2*, which encodes Ly-6-related protein (*43*), may contribute to neuronal hyperactivity independently from olfaction.

We found that aging causes hyperactivity of AWC sensory and AIA interneurons in *C. elegans*. Given that ablation of these neurons can restore behavior and that the effect of AWC hyperactivity is similar in young and aged animals, the age-dependent thermotaxis decline is not caused by loss of neurons or connectivity but by the change in neuronal excitability (Fig. 5). AWC sensory neurons and AIA interneurons can communicate through their bidirectional synapses, differentially mediated by neurotransmitter and neuropeptide signaling, to modulate the primary thermotaxis circuit of aged animals (Fig. 5). We show that in aged animals, AWC hyperactivity is dependent on *unc-13*, while AIA hyperactivity is dependent on *unc-31*. However, the remnant hyperactive signals of these neurons in respective mutants present the possible contribution of upstream neurons, likely shared by AWC and AIA. So, what is upstream of AWC and AIA? ASI neurons sense food cues to modulate diverse *C. elegans* behaviors (*44-46*) and make chemical synapses with AWC and AIA neurons and gap junctions with AIA interneurons (*34*). Moreover, ASI neurons play a role in regulating thermotaxis in young animals (*17*) (Fig. 2A). Compared to the young animals, our data revealed an opposite valence modulation of thermotaxis behavior in *E. coli*-fed aged animals upon ablation of ASI sensory neurons (Fig. 2A), suggesting an age-dependent regulation. Neuroendocrine signaling in ASI through neuropeptides (http://www.wormatlas.org) might aid in the crosstalk between AWC and AIA neurons age-dependently. Diets like *L. reuteri* might function in internal state-dependent pathways to ameliorate *C. elegans* age-dependent thermotaxis decline. In young animals, AWC displays starvation-dependent increased responses to thermal stimuli to impair thermotaxis (*25*). Furthermore, AWC increases inter-individual variation in AIY neuronal activity (*47*). These AWC properties in young animals might contribute to hyperactivation and behavioral impairment in aged animals. Previous studies on aged *C. elegans* involving other behavioral contexts have demonstrated that chronic excitation of oxygen-sensing URX neurons causes a loss of plasticity to oxygen concentration (*48*); also, increased activity of AVA interneuron, mediating backward locomotion, was seen to be consistent with increased tendency for reversals (*49*). In mice, hyperexcitable hypocretin neurons caused fragmented sleep during aging (*50*). Clinical studies in Alzheimer’s disease patients and studies with transgenic mice models of Alzheimer’s disease have shown hippocampal and cortical neuronal hyperactivity with abnormal levels of sodium, glutamate, and intracellular calcium (*51-56*). Furthermore, increased spontaneous activity, causing functional degradation, was found in the primary visual cortex of senescent monkeys (*57*). We suggest that neuronal hyperactivity may be a conserved mechanism that causes age-dependent cognitive decline.

**Fig. 5.**
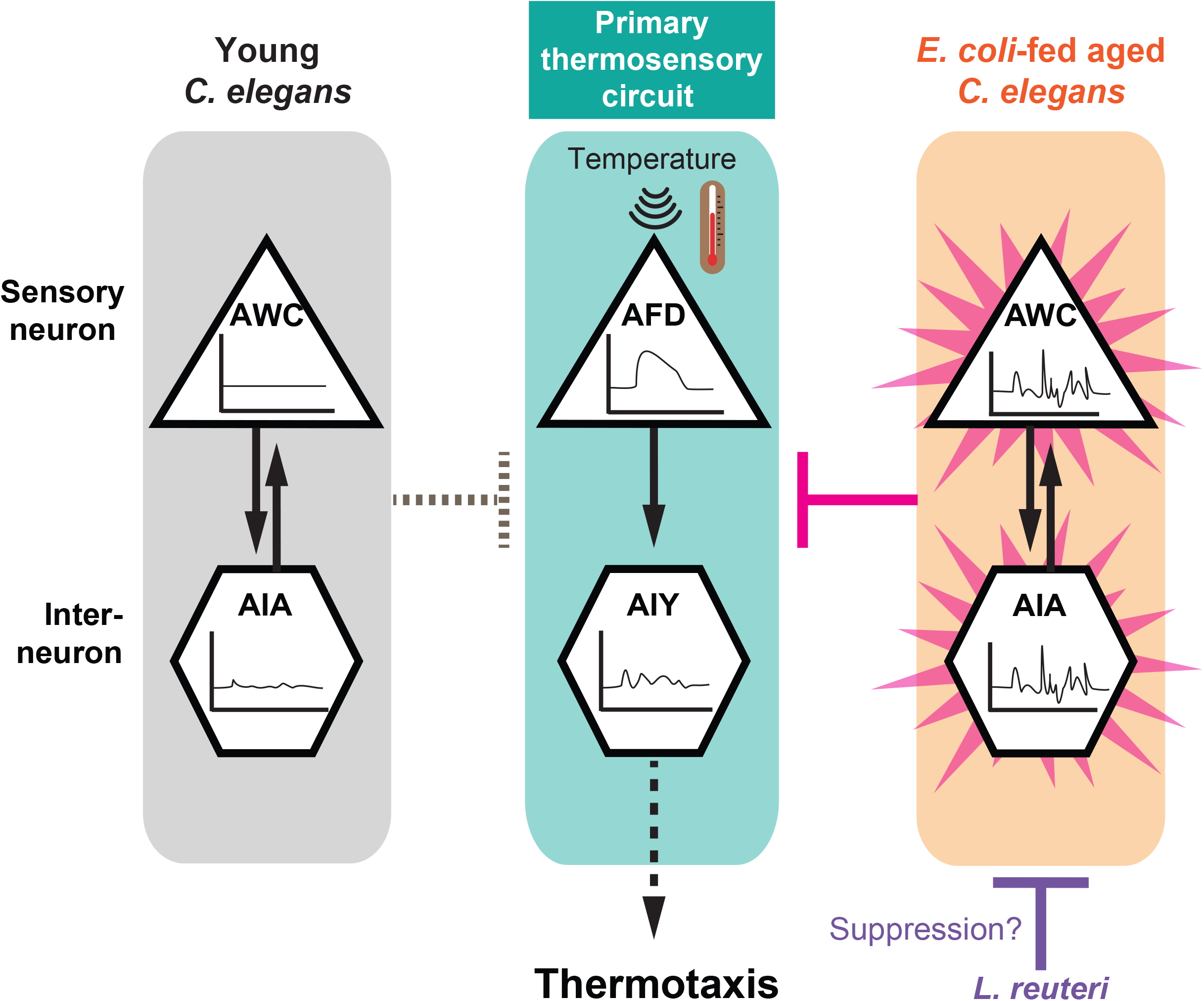
Model of age- and diet-dependent thermotaxis decline. AWC and AIA neurons, which are dispensable for thermotaxis in young animals, become hyperactive and interfere with the primary thermosensory circuit in aged animals. Feeding animals with *L. reuteri* ameliorates age-dependent thermotaxis decline by suppressing the AWC and AIA hyperactivity.

## Acknowledgments

We thank S. Nakano, H. J. Matsuyama, T. T. Huang, and A. Kano for their invaluable discussion and comments on this manuscript. We thank H. Bito (The University of Tokyo) for kindly providing the calcium indicator RCaMP2 plasmid. Some strains were provided by the CGC, which is funded by NIH Office of Research Infrastructure Programs (P40 OD010440).

## Author contributions

Conceptualization: BMA, MM, KN

Data curation: BMA, MM, AY, ST

Formal analysis: BMA, MM, MSJ

Funding acquisition: IM, KN

Investigation: BMA, MM, AY, ST

Methodology: BMA, MM, KN

Project administration: IM, KN

Resources: MSJ, YT, IM, KN

Software: BMA, MSJ, YT, IM

Supervision: IM, KN

Validation: BMA, KN

Visualization: BMA, MM, AY, ST, KN

Writing – original draft: BMA

Writing – review & editing: BMA, IM, KN

## Data and materials availability

All data are available in the main text or the supplementary materials.

## Supplementary Materials

Materials and Methods

Figs. S1 to S6

Table S1

References (*58-65*)

