## Supplementary Data for "Aberrant neuronal hyperactivation causes an age- and diet-dependent decline in associative learning behavior"

**The PDF file includes:**

Materials and Methods
Figs. S1 to S6
Tables S1
References (*58*-*65*)

#### Materials and Methods

C. elegans maintenance and strains

All *C. elegans* strains were maintained at 23°C on Nematode Growth Medium (NGM) plates with *Escherichia coli* (*E. coli*, OP50) as a food source, as previously reported (*58*). N2 (Bristol) was used as the wild type. Strains were obtained from the Caenorhabditis Genetic Center or generated in our laboratory using standard crossing or transgenesis methods as described below (*59, 60*). The strains used in this study are summarized in table S1.

Generation of plasmids and transgenic strains

The entry vectors based on pCR8 (Invitrogen) and destination vectors were cloned by Gibson assembly (NEB) and recombined using LR clonase (Invitrogen) to generate expression vectors. *gcy-8* promoter (*gcy-8*p) and *ges-1* promoter (*ges-1*p) contain ~0.7-kbp and ~3-kbp upstream sequence of the corresponding genes, respectively. The transgenic strain NUJ296 *knjIs15[gcy-8p::GCaMP6m + gcy-8p::TagRFP + ges-1p::TagRFP]* was generated by optogenetic mutagenesis (61). Briefly, the following plasmids were injected into CZ20310 *juSi164[mex-5p::his-72::miniSOG + Cb-unc-119] unc-119(ed3)* III, and animals were illuminated with blue light 48 hours after injection to induce integration: *gcy-8*p::GCaMP6m (pKEN1036) at 75 ng/µl, *gcy-8*p::TagRFP (pKEN1033) at 5 ng/µl, and *ges-1*p::TagRFP (pKEN848) at 20 ng/µl. The strain was crossed to N2 animals four times to remove *juSi164* and *unc-119(ed3)* and named NUJ296. The transgenic strain IK3824 *njEx2138[gcy-8p::NLS::RCaMP2 + AIYp::RCaMP2 + ges-1p::NLS::TagRFP]* was generated by injecting the following plasmids into N2: *gcy-8*p::NLS::RCaMP2 (pYT78) at 70 ng/µl, AIYp::RCaMP2 (pYT79) at 70 ng/µl, and *ges-1*p::NLS::TagRFP (pNAS88) at 70 ng/µl. RCaMP2 was a gift from Haruhiko Bito (*62*). For tissue-specific rescue experiments of *daf-16*, *daf-16* promoter of *daf-16bp::daf-16b* (pKEN1016) (*9*) was replaced with *gcy-8*p (pKEN1017), *ttx-3*p/AIYp (pKEN1018), *ceh-36*p (pKEN1054), and *ins-1*p (pKEN1055) using Gibson assembly. Using these plasmids, the single-copy inserted strains were generated as previously described (*63*) . Briefly, each plasmid was injected into N2 animals with the plasmid carrying Cas9+sgRNA for cxTi10882 site (pCZGY2750) and co-injection markers, kindly provided by Erik Jorgensen: *rab-3*p::mCherry (pGH8); *myo-2*p::mCherry (pCFJ90); *myo-3*p::mCherry (pCFJ104). Single-copy inserted animals were selected based on the hygromycin-resistance and the absence of any red signals from the co-injection markers.

Bacterial preparation

Bacterial handling was done using strict sterile techniques. *E. coli* (OP50) was inoculated into LB and cultured overnight at 37°C for regular worm maintenance. Concentrated bacteria were used for aged animals. *E. coli* was inoculated from a glycerol stock into Super Broth and cultured overnight at 37°C. *Lactobacillus reuteri* (also known as *Limosilactobacillus reuteri*) strain SBT10010 was provided by MEGMILK SNOW BRAND company and deposited in the International Patent Organism Depositary, National Institute of Technology and Evaluation (Chiba, Japan) under the accession number NITE P-02997. *L. reuteri* was inoculated into MRS broth (Becton Dickinson Co.) from a glycerol stock, cultured, and passaged a few times. The 1/100 volume of bacterial solution was inoculated into Super Broth for *E. coli* or MRS for *L. reuteri*, and cultured 16 hours at 37°C without shaking. Bacterial cells were collected by centrifugation for 3 (*E. coli*) and 10 min (*L. reuteri*) at 7000 x g and 4°C. Subsequently, cells were washed twice with 0.9% NaCl and adjusted to a final concentration of 0.1 g/ml (wet weight) with NG buffer (25 mM

K-$PO_4$ (pH 6), 50 mM NaCl, 1 mM $CaCl_2$, 1 mM $MgSO_4$). Two hundred microliters of the reconstituted bacterial suspension were spread onto 60 mm NGM plates and dried overnight.

Preparation of animals for behavioral assays and imaging

Animals for behavioral assays were prepared by bleaching gravid hermaphrodites using a 1:1 mixture of household bleach and 1M NaOH. The resulting synchronized eggs were placed onto *E. coli*-seeded NGM plates and cultivated at 23°C for 72 hr to obtain day-one adults (D1, young). For thermotaxis of aged animals, D1 animals were daily washed with NG buffer and transferred to fresh NGM plates seeded with 0.1 g/ml *E. coli* or *L. reuteri* for four days until they reached day-five adults (D5, aged). For calcium imaging, three gravid hermaphrodites were picked and allowed to lay eggs on an *E. coli*-seeded NGM plate. After 48 hr, 30-40 L4 larvae with the transgene were selected onto a fresh NGM plate under a fluorescence microscope and cultivated for another 24 hr to obtain D1. To obtain aged animals, 30-40 D1 animals were picked onto respective bacterial plates daily. To synchronize animals for imaging, L4 animals with transgenes were selected. The following day, the selected animals which had become D1 adults were imaged as young animals. To image aged animals, D1 adults were transferred by a worm pick onto *E. coli*- or *L. reuteri*-seeded NGM plates every day until reaching D5.

Thermotaxis assay

Fifty to 250 animals were used for population thermotaxis assays on a linear thermal gradient as previously reported (*7*). Animals were washed off cultivation plates with NG buffer and placed at the center of assay plates (2% w/v Bacto Agar, 0.3% w/v NaCl, 25 mM K-$PO_4$ (pH 6)) with a 17-23°C temperature gradient at 0.45-0.5°C/cm without food. The one-hour assay was stopped by killing animals by applying chloroform on the lid, and the number of worms in each of the eight sections of the plate was counted. Thermotaxis ability was evaluated by calculating TTX performance index (Fig. 1C), TTX index (fig. S2A), and Normalized TTX performance index (fig. S4). For thermotaxis of mito-miniSOG cell-ablated strains (*15, 24*), L4 larvae and D4 adult animals on food-containing plates were continuously exposed to blue light (488 nm) for 7.5 min without lids for D1 and D5 assays, respectively, 24 hr prior to the assay. The intensity of blue light received by the animals was measured as 2 $mW/mm^2$. For thermotaxis of *daf-16* rescue strains (fig. S4), inter-genotype variations of same age and diet groups was corrected by subtracting the mean TTX performance index of respective *E. coli*-fed aged group and then normalized by dividing by the mean TTX performance index of the respective D1 group.

Calcium imaging

Calcium imaging was performed on immobilized animals as previously described (*64*). Animals expressed the following genetically encoded calcium sensors in neuron(s) of interest: RCaMP2 in AFD and AIY, GCaMP6m and TagRFP in AFD, GCaMP6f and TagRFP in AWC, or YCX1.6 in AIA. Animals were picked off food and transferred to a mixture of 1 µL of 50-nm polystyrene microbeads (Polysciences) and 1 µL of 10 mM levamisole (Sigma) on a ~1 mm thick 10% agarose pad mounted on a 24-mm square cover glass. A circular cover slip of 15-mm diameter was placed atop the solution to immobilize the animals. The samples were then placed on a Peltier-based temperature controller set at the starting temperature. Samples were left for ~3 min before applying thermal stimulus. For AFD single imaging, we applied a thermal stimulus of 19-24°C; pausing at 19°C for 20 sec, rising linearly at 0.02°C/s to 24°C, and then pausing at 24°C for 180 sec. For AFD-AIY simultaneous and AWC imaging, we applied a thermal stimulus of 21.5-22.5°C;

pausing at 21.5°C for 30 sec, rising linearly at 0.05°C/s to 22.5°C, pausing at 22.5°C for 30 sec, and then falling back to 21.5°C. For AWC and AIA temperature-dependency assays, a constant thermal stimulus of 21.5°C was delivered to the animals throughout the duration of the assay (120 sec). Epifluorescent images were acquired with the 60X objective (NA = 0.9) of the upright BX61WI microscope (Olympus). One image was taken every second with 400 ms (100 ms for AIA imaging) exposure time. Recordings were taken twice per strain on different days except for those in *unc-13(s69)* and *unc-31(e928)* backgrounds which had to be carried out more than twice to obtain appreciable sample sizes.

Calcium imaging analysis

Analysis of fluorescence intensities for GCaMP or FRET for YCX was performed using MetaMorph software (Molecular Devices), while that for AFD-AIY simultaneous imaging was done using ImageJ (NIH). For AFD and AIY simultaneous imaging (Fig. 1I-K), calcium activity, $F(t)/F_{min}$, was evaluated by normalizing the RCaMP2 fluorescence of each neuron with the minimum fluorescence value in each recording. For other neuronal cell imaging, calcium activity $R(t)/R_{min}$, was evaluated by taking the GCaMPf/TagRFP, GCaM6m/TagRFP or YFP/CFP fluorescence ratio, R(t), which was then standardized within each recording by normalizing with the minimum fluorescence ratio, $R_{min}$. AFD responses were quantified as "AFD onset temperature" defined as the temperature at or closest to half maximum AFD fluorescence value, AFD [$F_{max}$], or half maximum AFD fluorescence ratio, AFD [$R_{max}$] (Fig. 1L; fig S1E) (*8*). AIY response was quantified as the AIY fluorescence value, AIY [F], corresponding to AFD [$F_{max}$]. Spontaneous AWC and AIA activities were quantified by calcium-spike deconvolution and denoising using a modified custom-written MATLAB (Mathworks) code (*65*). The spike frequency and spike area per soma were then plotted for all conditions. Heat maps and activity plots were generated on MATLAB (Mathworks).

Statistical analysis

Graphs were generated using RStudio (https://www.r-project.org), and all data were analyzed for statistics using R (version 4.3.1). In box and whisker plots, the lower boundary, median line, and upper boundary of the boxes indicate the 25th, 50th, and 75th percentiles, while the whiskers show the minimum and maximum values. Each data point for thermotaxis assay is obtained from a single plate. In violin plots, middle lines within violins show medians, and top and bottom lines show interquartile ranges. For all experiments, Kruskal Wallis test was used to analyze statistical differences, with post-hoc Steel-Dwass tests performed for age- and diet- comparisons within a genotype. Post-hoc Steel test was used for corresponding comparisons with control within a condition. As an exception, for the thermophilicity experiment (fig. S2A), post-hoc Steel test was performed between D1 wild type and all other conditions. Different alphabets show significant difference. Post-hoc Steel test was performed for corresponding comparisons to control. $^{*}p \leq 0.05$, $^{**}p \leq 0.01$, and $^{***}$ $p \leq 0.001$, and 'ns' indicates not significant, $p>0.05$.

### Figure S1

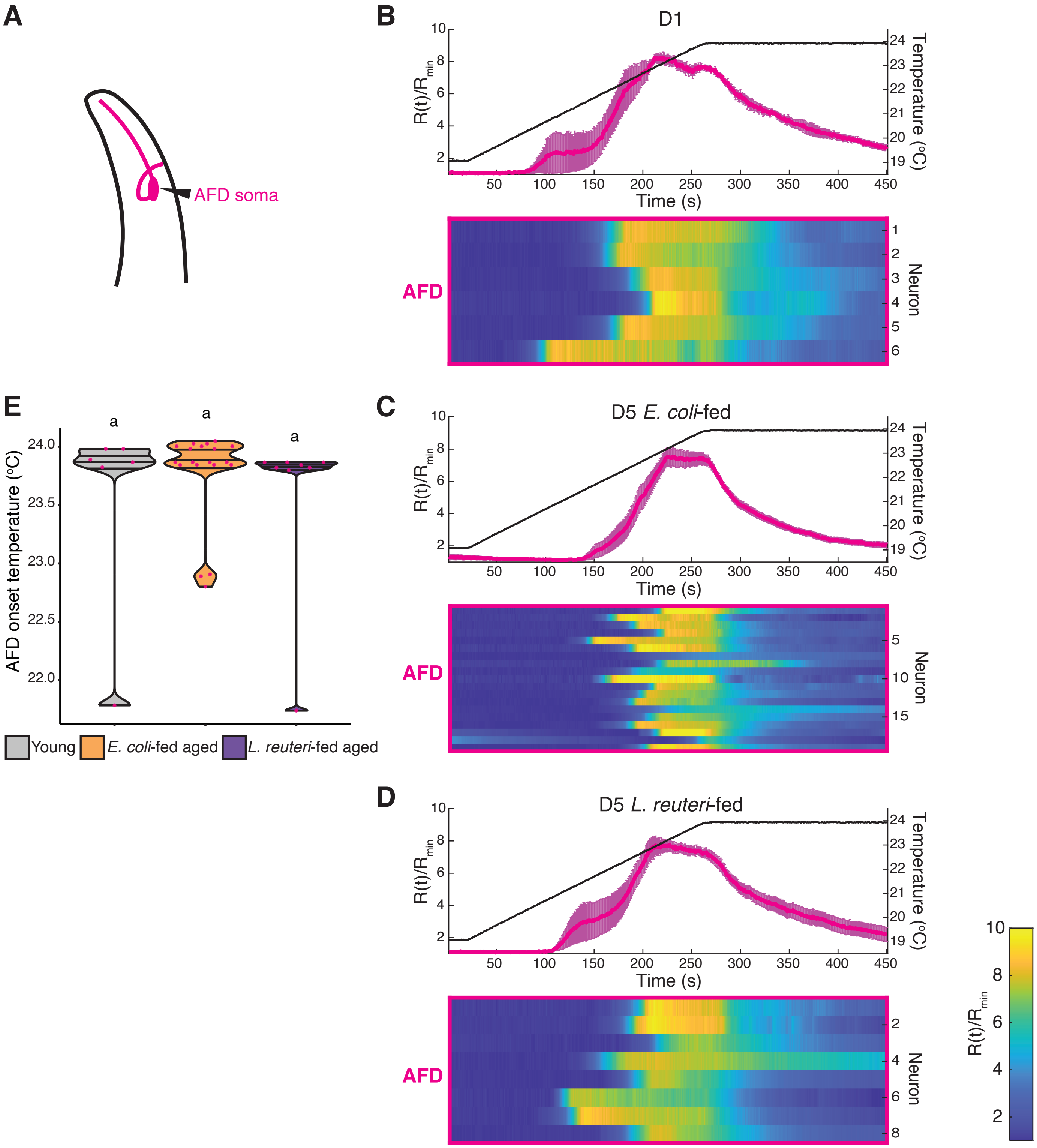

A
AFD soma
B
D1
C
D5 E. coli-fed
D
D5 L. reuteri-fed
R(t)/R_min
Temperature (°C)
Time (s)
AFD
Neuron
E
AFD onset temperature (°C)
a
Young
E. coli-fed aged
L. reuteri-fed aged

**Fig. S1.**

**Temperature-dependent AFD calcium activities are similar in young and aged animals irrespective of diets.**

**(A)** Schematic of AFD thermosensory neuron.
**(B,C,D)** Calcium signals of AFD soma in indicated conditions in the wild type in response to a linear temperature increase (5°C) at 0.02°C/s. GCaMP6m and TagRFP (used as a reference) in AFD were measured. The GCaMP6m/TagRFP ratio, R(t), was calculated and standardized by the minimum fluorescence ratio, $R_{min}$, in each neuron recording. Magenta lines depict the average calcium signals in AFD. Error bars are SEM. Black lines indicate average temperature stimuli. Heat maps of calcium signals are shown at the bottom.
**(E)** Violin plots showing temperature of AFD activation onset in from data in (B,C,D). Statistical analysis was by Kruskal Wallis test followed by post-hoc Steel-Dwass test for comparisons within a genotype. Same alphabets indicate no significant difference.

### Figure S2

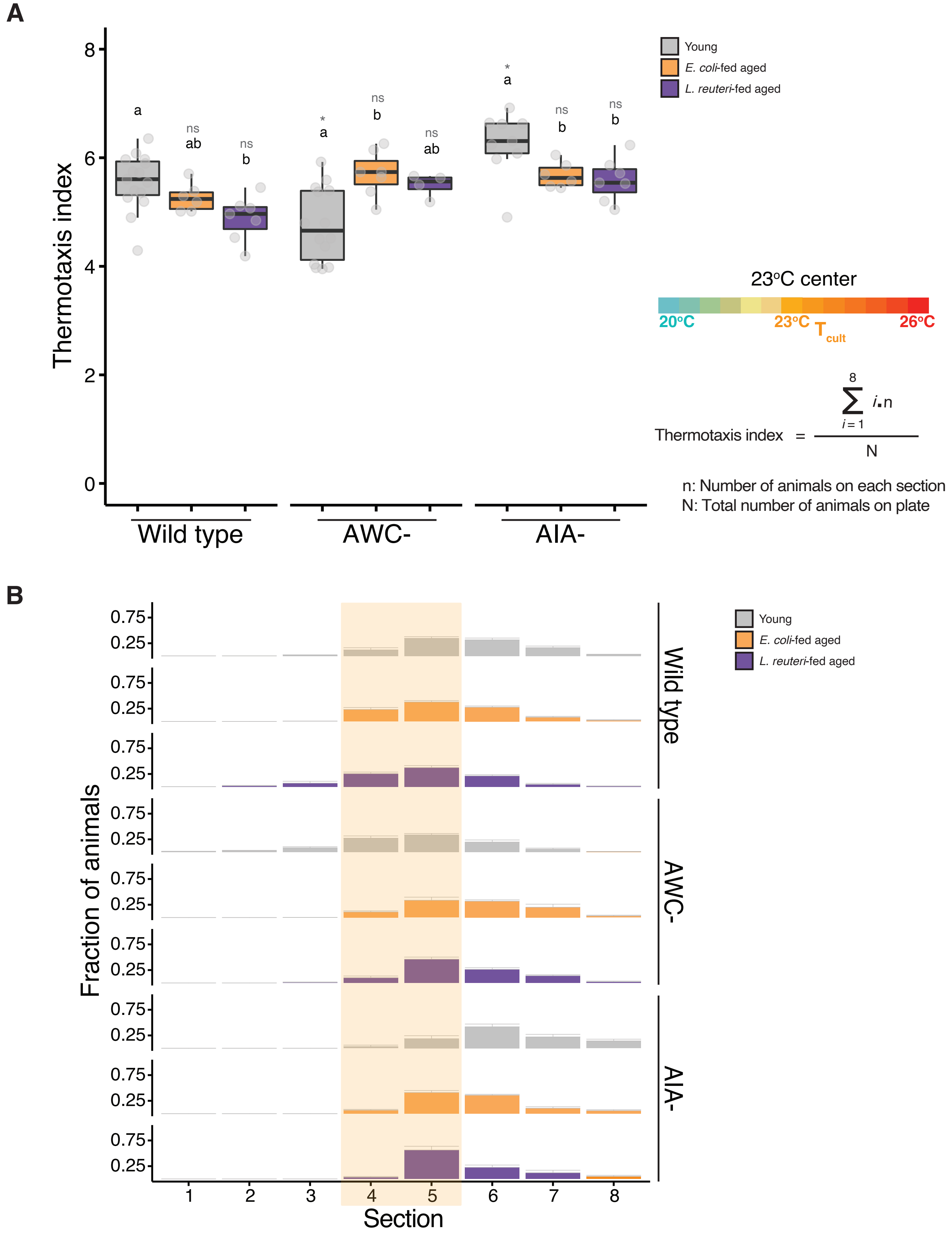

A
Thermotaxis index
8
6
4
2
0
a
ns
ab
ns
b
*
a
ns
b
ns
ab
*
a
ns
b
ns
b
Wild type
AWC-
AIA-
Young
E. coli-fed aged
L. reuteri-fed aged
23°C center
20°C
23°C
$T_{cult}$
26°C
Thermotaxis index = $\frac{\sum_{i=1}^{8} i \cdot n}{N}$
n: Number of animals on each section
N: Total number of animals on plate
B
Fraction of animals
0.75
0.25
Wild type
AWC-
AIA-
1
2
3
4
5
6
7
8
Section

**Fig. S2.**
**AWC- and AIA-ablated animals are not thermophilic.**

**(A)** Box and whisker plots of thermotaxis indices of wild type, AWC- and AIA-ablated animals. Animals cultivated at 23°C were spotted at a 23°C-center temperature gradient to test for thermophilicity.
**(B)** Distribution of animals with indicated genotypes on thermotaxis plates from data in (A). Light brown rectangles indicate the two sections near $T_{cult}$. Error bars denote standard error. Statistical analysis was by Kruskal Wallis test followed by post-hoc Steel-Dwass test for comparisons within a genotype. Different alphabets depict significant difference. Post-hoc Steel test was used for comparisons with young wild type animals only. *$p \leq 0.05$ and 'ns' indicates not significant ($p>0.05$).

#### Figure S3

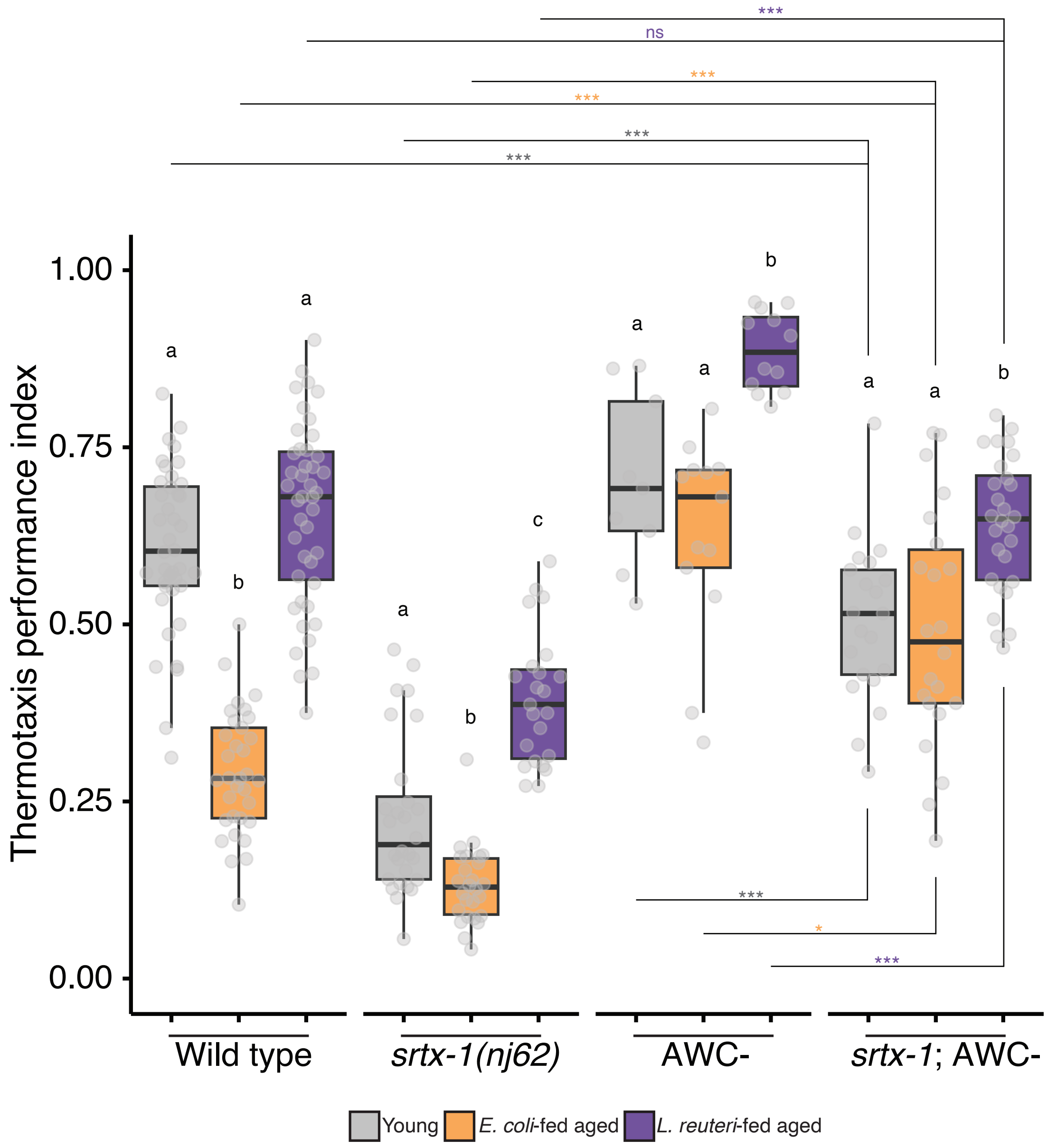
***
ns
***
***
***
***
Thermotaxis performance index
1.00
0.75
0.50
0.25
0.00
a
b
a
a
b
c
a
a
b
a
a
b
***
*
***
Wild type
srtx-1(nj62)
AWC-
srtx-1; AWC-
Young
E. coli-fed aged
L. reuteri-fed aged

**Fig. S3.**
**Thermotaxis defects of *srtx-1* mutants partially act through AWC.**

Box and whisker plots of thermotaxis performance indices of the wild type and indicated genotypes in the different diet and age conditions. *srtx-1(nj62)* mutants were used as a model of AWC hyperactivity, and AWC was ablated by the expression of caspase.
Statistical analysis was by Kruskal Wallis test followed by post-hoc Steel-Dwass test for comparisons within a genotype. Different alphabets depict significant difference. Post-hoc Steel test was used for corresponding comparisons with control within a condition, as represented by the distinct asterisk colors. *$p \leq 0.05$, ***$p \leq 0.001$, and 'ns' indicates not significant ($p>0.05$).

**Figure S4**

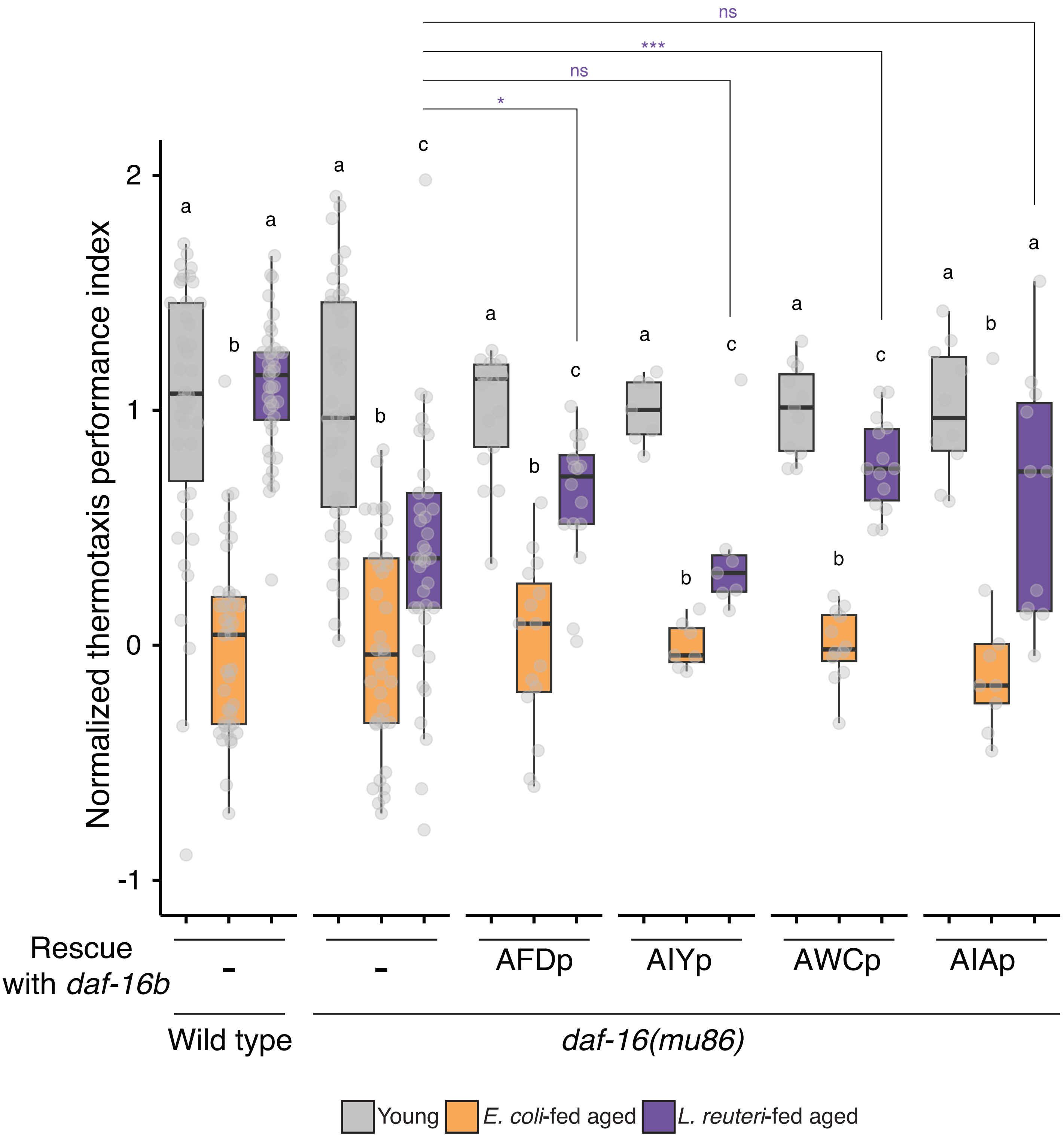
ns
***
ns
*
a
b
a
a
b
c
a
b
c
a
b
c
a
b
c
a
b
a
2
1
0
-1
Normalized thermotaxis performance index
Rescue with daf-16b
-
-
AFDp
AIYp
AWCp
AIAp
Wild type
daf-16(mu86)
Young
E. coli-fed aged
L. reuteri-fed aged

**Fig. S4.**

***daf-16* functions in AWC to ameliorate the age-dependent thermotaxis decline in *L. reuteri*-fed aged animals.**

Box and whisker plots of thermotaxis performance indices of wild type and indicated genotypes in the different diet and age conditions. The b isomer of *daf-16* was used to rescue *daf-16(mu86)* mutant phenotype in indicated neurons.

Statistical analysis was by Kruskal Wallis test followed by post-hoc Steel-Dwass test for comparisons within a genotype. Different alphabets depict significant difference. Post-hoc Steel test was used for corresponding comparisons with control within a condition, as represented by the distinct asterisk colors. *$p \leq 0.05$, ***$p \leq 0.001$, and 'ns' indicates not significant ($p>0.05$).

#### Figure S5

**A**

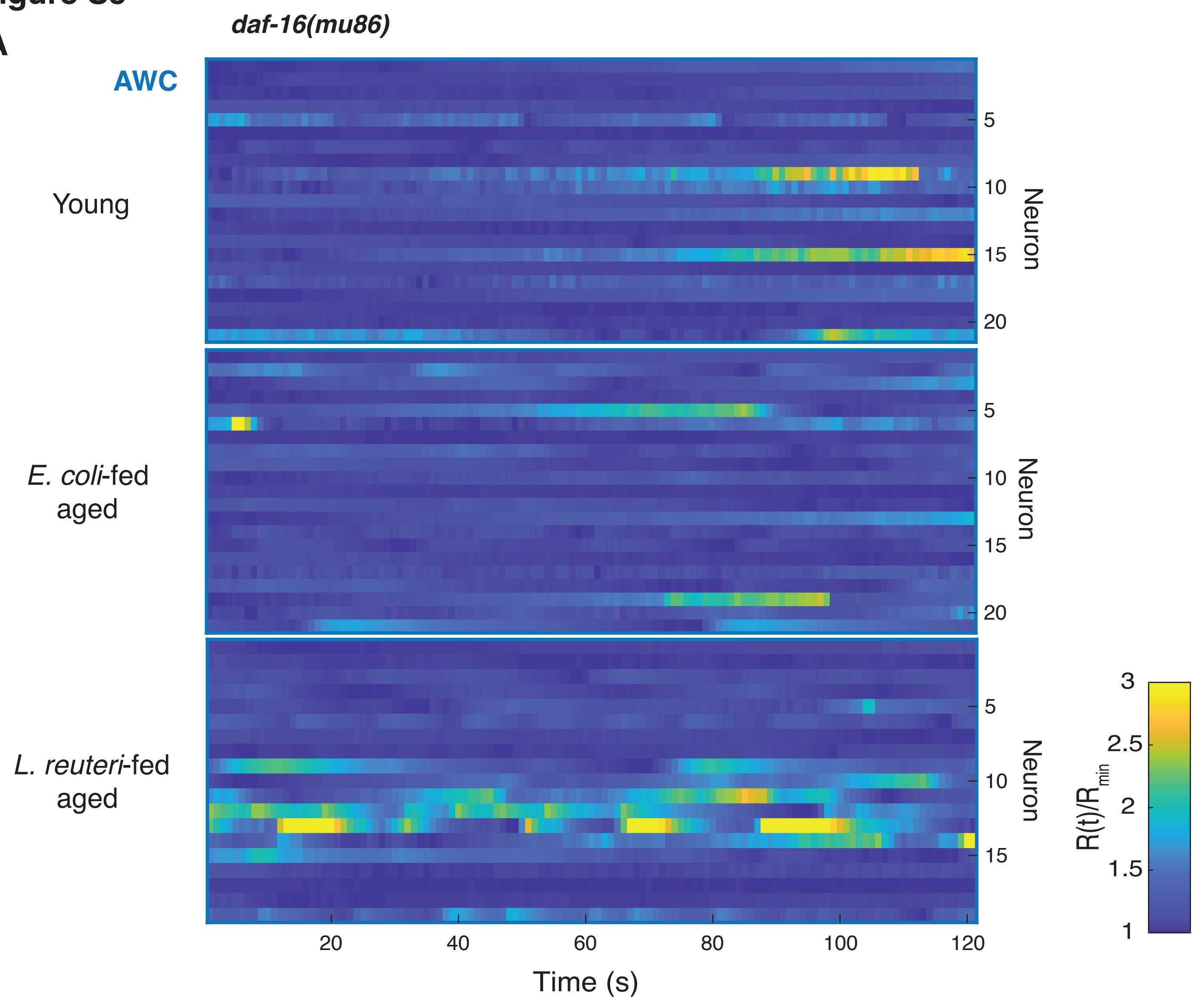
daf-16(mu86)
AWC
Young
E. coli-fed aged
L. reuteri-fed aged
Neuron
5
10
15
20
20
40
60
80
100
120
Time (s)
R(t)/R_min
3
2.5
2
1.5
1

**B**

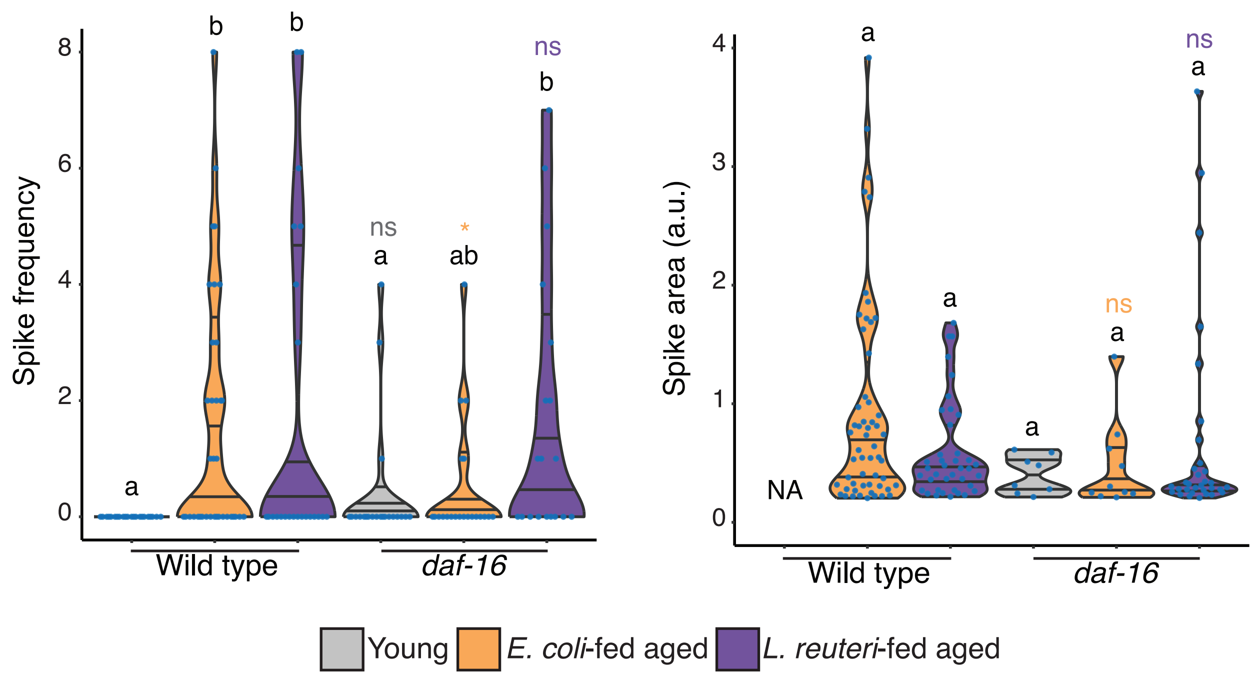
Spike frequency
Spike area (a.u.)
Wild type
daf-16
a
b
ab
ns
*
NA
Young
E. coli-fed aged
L. reuteri-fed aged

**Fig. S5.**
***daf-16* mutants show increased hyperactivity of *L. reuteri*-fed aged animals.**

**(A)** Heat maps of calcium signals of AWC soma in indicated conditions in *daf-16(mu86)* mutants, which failed to show amelioration of age-dependent thermotaxis decline with *L. reuteri* feeding, under a constant temperature stimulus of 21.5$^{o}$C.
**(B)** Quantification of calcium spike frequency per soma (left) and calcium spike area per soma (right) from data in (A). Wild type data are the same as in Fig. 2K.

### Figure S6

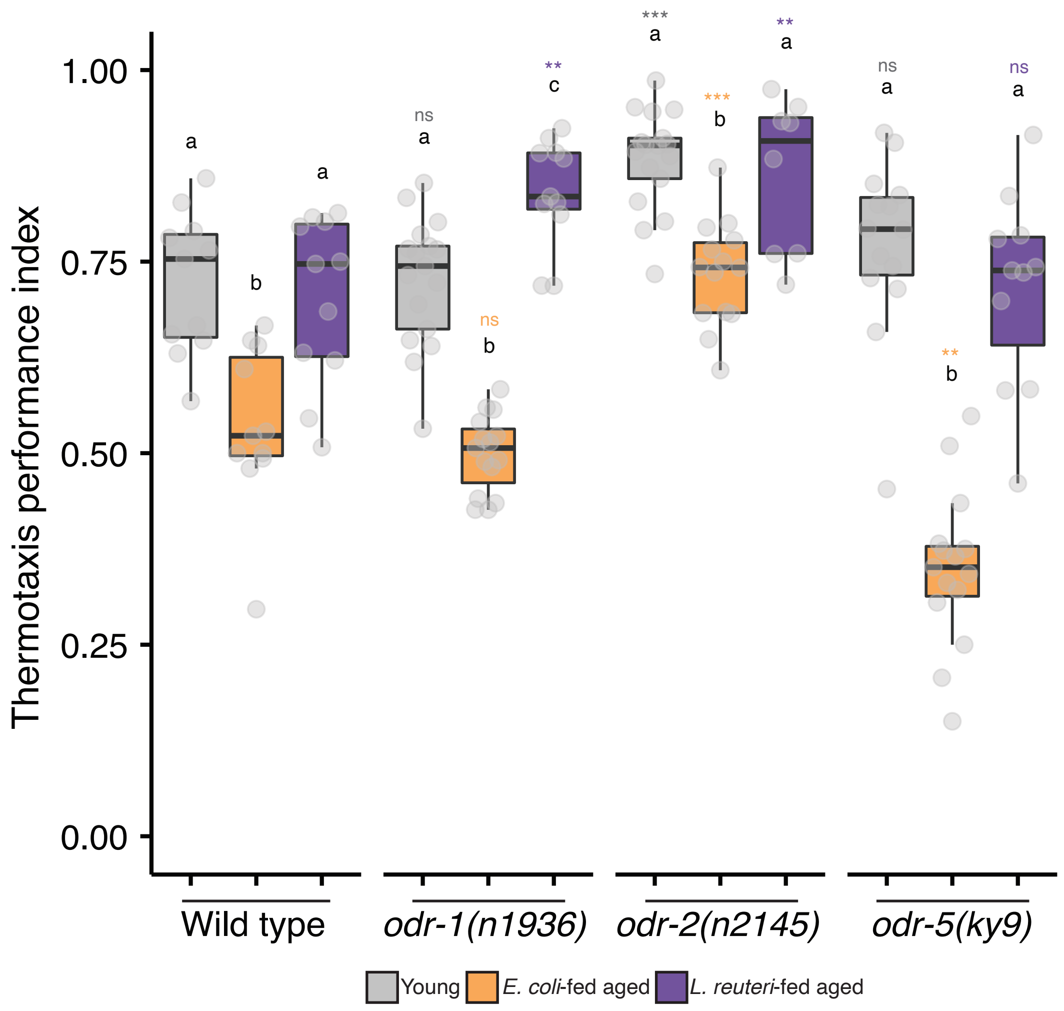

Thermotaxis performance index
1.00
0.75
0.50
0.25
0.00
a
b
a
ns
a
ns
b
**
c
***
a
***
b
**
a
ns
a
**
b
ns
a
Wild type
*odr-1(n1936)*
*odr-2(n2145)*
*odr-5(ky9)*
Young
*E. coli*-fed aged
*L. reuteri*-fed aged

**Fig. S6.**
**AWC and AIA hyperactivity is not acting through olfaction.**

Box and whisker plots of thermotaxis indices of the wild type and indicated mutants defective in chemotaxis behavior toward AWC-sensed odorants.
Statistical analysis was by Kruskal Wallis test followed by post-hoc Steel-Dwass test for comparisons within a genotype. Different alphabets depict significant difference. Post-hoc Steel test was used for corresponding comparisons with control within a condition, as represented by the distinct asterisk colors. *$p \leq 0.05$, *$p \leq 0.01$, *$p \leq 0.001$, and 'ns' indicates not significant ($p>0.05$).

**Table S1 *C. elegans* strains used in the study**

| Strain | Genotype | Feature / Purpose |
|---|---|---|
| N2 | N2 Bristol | *C. elegans* wild type |
| IK2809 | *njIs80[gcy-8p::cz::casp3(p17) + gcy-8p::casp3(p12)::nz + ges-1p::NLS::GFP] X* | AFD ablation |
| IK3048 | *njIs89[gcy-8p::tomm-20(N'-55AA)::miniSOG + ges-1p::GFP] III* | AFD ablation |
| IK2710 | *njIs62[AIYp::cz::casp3(p17) + AIYp::casp3(p12)::nz + ges-1p::NLS::GFP] V* | AIY ablation |
| IK2808 | *njIs79[ceh-36p::cz::casp3(p17) + ceh-36p::casp3(p12)::nz + ges-1p::NLS::GFP] X* | AWC ablation |
| IK3125 | *njIs98[ceh-36p::tomm-20(N'55AA)::miniSOG + ges-1p::NLS::GFP] I* | AWC ablation |
| PY7505 | *oyIs84[gcy-27p::cz::casp3(p17) + gpa-4p::casp3(p12)::nz + gcy-27p::GFP + unc-122p::dsRed]* | ASI ablation |
| IK3179 | *njIs107[acc-2p::cz::casp3(p17) + odr-2(2b)p::casp3(p12)::nz + ges-1p::NLS::GFP] IV* | AIZ ablation |
| IK3449 | *njIs133[aptf-1p::cz::casp3(p17) + inx-1p::casp3(p12)::nz + ges-1p::NLS::GFP] II* | AIB ablation |
| IK3263 | *njIs120[ins-1p::cz::casp3(p17) + gcy-28dp::casp3(p12)::nz + ges-1p::NLS::GFP] V* | AIA ablation |
| IK3240 | *njIs115[ins-1p::FLP + gcy-28dp::FRT::tomm-20(N'-55AA)::miniSOG + ges-1p::NLS::GFP] IV* | AIA ablation |
| IK2910 | *njIs84[glr-3p::cz::casp(p17) + glr-3p::casp(p12)::nz + ges-1p::NLS::GFP] III* | RIA ablation |
| NUJ296 | *knjIs15[gcy-8Mp::GCaMP6m + gcy-8Mp::TagRFP + ges-1p::TagRFP]* | For AFD calcium imaging |
| IK3824 | *njEx2138[gcy-8p::NLS::RCaMP2 + AIYp::RCaMP2 + ges-1p::NLS::TagRFP]* | For AFD and AIY simultaneous calcium imaging |
| IK3296 | *njEx1368[ceh-36p::GCaMP6f + str-2p::TagRFP]* | For AWC calcium imaging |
| NUJ654 | *unc-13(s69) I; njEx1368[ceh-36p::GCaMP6f + str-2p::TagRFP]* | For AWC calcium imaging in mutant |
| NUJ655 | *unc-31(e928) IV; njEx1368[ceh-36p::GCaMP6f + str-2p::TagRFP]* | For AWC calcium imaging in mutant |
| NUJ656 | *daf-16(mu86) I; njEx1368[ceh-36p::GCaMP6f + str-2p::TagRFP]* | For AWC calcium imaging in mutant |
| NUJ556 | *njIs120[ins-1p::cz::casp3(p17) + gcy-28dp::casp3(p12)::nz + ges-1p::NLS::GFP] V; njEx1368[ceh-36p::GCaMP6f + str-2p::TagRFP]* | For AWC calcium imaging in AIA-ablated background |
| IK3331 | *njEx1387[ins-1p::FLP + gcy-28dp::FRT::stop::FRT::stop::YCX1.6]* | For AIA imaging |
| NUJ594 | *unc-13(s69) I; njEx1387[ins-1p::FLP + gcy-28dp::FRT::stop::FRT::stop::YCX1.6]* | For AIA imaging in mutant |
| NUJ593 | *unc-31(e928) IV; njEx1387[ins-1p::FLP + gcy-28dp::FRT::stop::FRT::stop::YCX1.6]* | For AIA imaging in mutant |
| NUJ572 | *njIs79[ceh-36p::cz::casp3(p17) + ceh-36p::casp3(p12)::nz + ges-1p::NLS::GFP] X; njEx1387[ins-1p::FLP + gcy-28dp::FRT::stop::FRT::stop::YCX1.6]* | For AIA calcium imaging in AWC-ablated background |
| IK646 | *srtx-1(nj62) IV bc11* | Mutant |
| NUJ601 | *srtx-1(nj62) IV; njIs79[ceh-36p::cz::casp3 + ceh-36p::casp3::nz + ges-1p::GFP]* | Mutant |
| CF1038 | *daf-16(mu86) I* | Mutant |
| NUJ353 | *daf-16(mu86) I; knjSi20[gcy-8p::daf-16b] IV* | For AFD-specific *daf-16* rescue |
| NUJ427 | *daf-16(mu86) I; knjSi28[AIYp::daf-16b] IV* | For AIY-specific *daf-16* rescue |
| NUJ509 | *daf-16(mu86) I; knjSi29[ceh-36p::daf-16b] IV* | For AWC-specific *daf-16* rescue |
| NUJ537 | *daf-16(mu86) I; knjSi30[ins-1p::daf-16b] IV* | For AIA-specific *daf-16* rescue |
| CX2065 | *odr-1(n1936) X* | Mutant |
| CX2304 | *odr-2(n2145) V* | Mutant |
| CX2357 | *odr-5(ky9) X* | Mutant |
